# Clonal Heterogeneity Supports Mitochondrial Metabolism in Pancreatic Cancer

**DOI:** 10.1101/2020.05.15.098368

**Authors:** Christopher J. Halbrook, Galloway Thurston, Amy McCarthy, Barbara S. Nelson, Peter Sajjakulnukit, Abigail S. Krall, Peter J. Mullen, Li Zhang, Sandeep Batra, Andrea Viale, Ben Z. Stanger, Heather R. Christofk, Ji Zhang, Marina Pasca di Magliano, Claus Jorgensen, Costas A. Lyssiotis

## Abstract

Pancreatic ductal adenocarcinoma (PDA) is characterized by a heterogenous and densely fibrotic microenvironment. This limits functional vasculature and diffusion of nutrients through the tumor^1,2^. Accordingly, pancreatic cancer cells develop numerous metabolic adaptations to survive and proliferate in nutrient austere conditions^3-7^. Subtypes of PDA have been characterized by transcriptional and functional differences^8-12^, which have been reported to exist within the same tumor^13-15^. However, it remains unclear if this diversity extends to metabolic programming. Here, using a combination of metabolomic profiling and functional interrogation of metabolic dependencies, we identify two distinct metabolic subclasses within neoplastic populations isolated from a single pancreatic tumor. Furthermore, these populations are poised for metabolic crosstalk, and in examining this, we find an unexpected role for asparagine in maintaining cell proliferation following mitochondrial inhibition. Functionally, when challenged by mitochondrial inhibition, asparagine supplementation increases intracellular levels of asparagine and aspartate, a rate limiting biosynthetic precursor^16-18^. Conversely, depletion of extracellular asparagine with PEG-asparaginase sensitizes pancreatic tumors to mitochondrial targeting with phenformin. Together, these data extend the concept of metabolic diversity to neoplastic populations within individual tumors, while illustrating a new method of intratumoral communication that supports tumor fitness^19,20^. Finally, the combination of asparaginase with mitochondrial inhibition could provide a powerful new strategy for this difficult to treat disease.

## Introduction

The tumor microenvironment in pancreatic ductal adenocarcinoma (PDA) is a complex ecosystem^21^, with diverse populations of fibroblasts and immune cells functioning to create a niche supporting cancer cell survival and tumor growth^19,20,22-28^. Accordingly, this allows for many and varied cooperative intratumoral crosstalk interactions in the microenvironment^29,30^. These extend to direct metabolic support of cancer cells from adjacent fibroblasts and tumor associated macrophages^31-35^.

The majority of PDAs express mutant Kras, so early efforts to understand metabolism in pancreatic cancer focused on the cell intrinsic metabolic rewiring downstream of Kras signaling pathways^36-38^. In addition, extensive metabolic profiling of a large set of PDA cell lines revealed heterogeneity among preferred bioenergetic pathways and sensitivity to metabolic inhibitors^12^. However, given the diversity of the neoplastic cell populations found in pancreatic tumors^19,20^, we postulated clonal differences within the cancer cells might also extend to metabolic behaviors, capable of symbiotic support^39^.

### Metabolic Characterization Stratified PDA Clones into Two Classes

To identify clonal differences in metabolism, we profiled a series of clonal cell lines derived from a single *Kras*^+/G12D^;*Trp53*^+/R172H^;*Pdx1*-Cre (KPC) mouse tumor (**Fig. 1A**) by liquid chromatography-coupled tandem mass spectrometry (LC/MS)-based metabolomics. Through unbiased clustering analysis, we observed that the clones clearly separated into two distinct groups based on their steady state metabolite pools (**Fig. 1B**). Pathway enrichment analysis demonstrated that these differences extended across several pathways, with glycolysis and Warburg effect metabolism topping the list (**Extended Data Fig. 1**). Indeed, we find that clones V, E, and H (hereby termed group 1) demonstrated increased levels of glycolytic metabolites (**Fig. 1C**) as compared to those found in clones K, M, N, and T (termed group 2). This difference in glycolysis between groups was confirmed by measuring lactate production in conditioned media of each of the clones (**Fig. 1D**).

**Figure 1:**
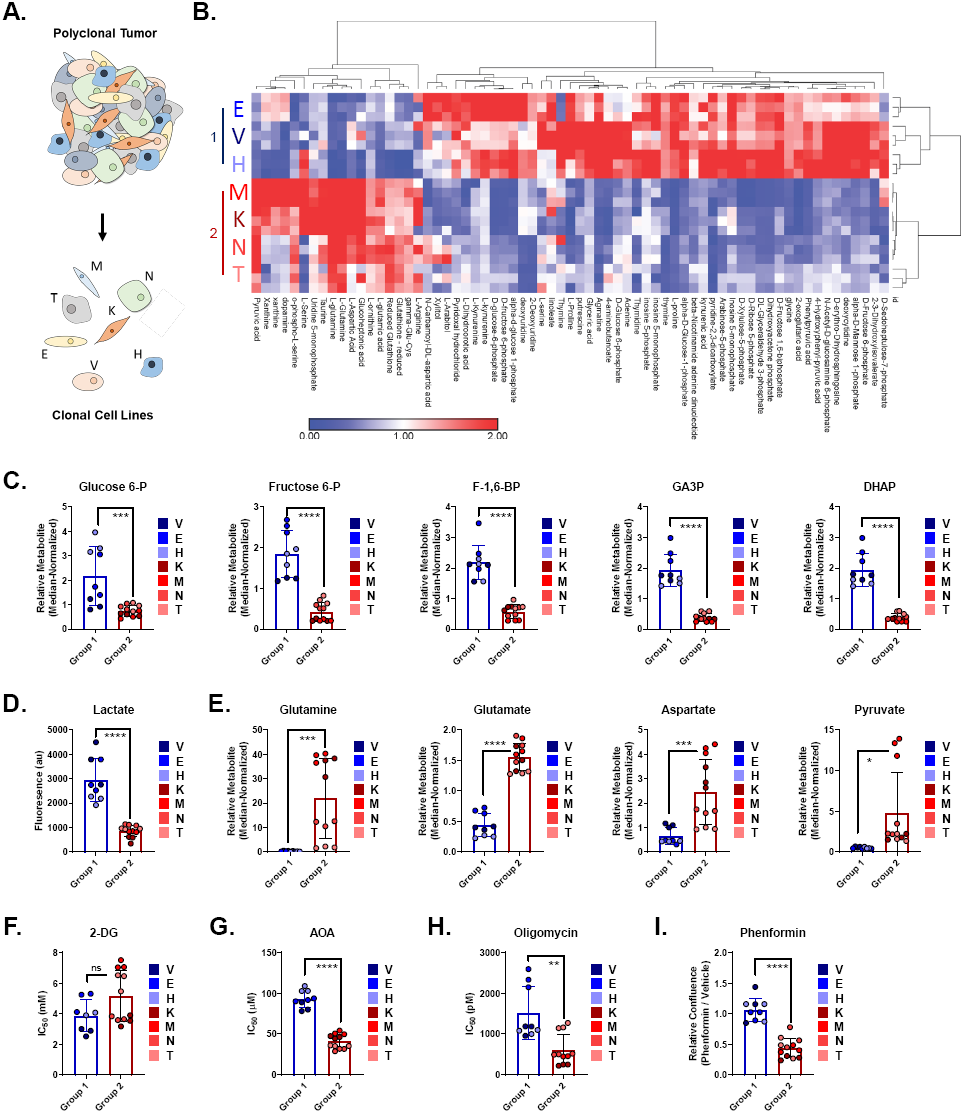
Characterization of Clonal Metabolites Reveals Two Distinct Populations. **A.** A polyclonal cell line established from a murine pancreatic tumor was subcloned into 7 clonal cell lines. **B**. Heatmap representation of significant intercellular metabolites present in the clonal cell lines grown in the same media conditions ±2 fold difference, *P* = 0.01 between group 1 clones (E,V,H) vs group 2 (K,M,N,T). Rows are clonal cells lines in triplicate and columns represent metabolites (n=3 replicates per cell line). **C**. Relative abundance of metabolites from the glycolysis pathway across clonal populations (n=3 replicates per cell line). **D.** Lactate production of clonal cell lines (n=3 replicates per cell line). **E.** Relative abundance of selected glutamine and TCA cycle branching metabolites (n=3 replicates per cell line). **F-I**. Sensitivity of clonal cell lines to 2-deoxyglucose (2-DG) (**F**), aminooxyacetic acid (AOA) (**G**), oligomycin (**H**), presented as the IC_50_, or for 25 µM phenformin (**I**) (n=3 replicates per cell line). F-1,6-BP = fructose 1,6-bisphsophate, GA3P = glyceraldehyde-3-phosphate, DHAP = dihydroxyacetone phosphate. Error bars are mean ±SD, * *P* ≤ 0.05; ** *P* ≤ 0.01; *** *P* ≤ 0.001; **** *P* ≤ 0.0001.

In contrast, we observed that group 2 clones demonstrated increased levels of glutamine and glutamate relative to group 1 (**Fig. 1E**). This observation was intriguing, as we have previously found that PDA cells utilize glutamine derived carbon to fuel TCA metabolism, which suggested that these clones may exhibit differential dependency on anabolic mitochondrial programs. Indeed, we also observed increased levels of pyruvate and aspartate in these group 2 clones relative to group 1.

Given that the clones separated into two groups based on basal metabolism, we next sought to determine if there were also differential metabolic vulnerabilities between these groups. While we did not observe a difference in sensitivity to inhibition of glycolysis through treatment with 2-deoxyglucose (**Fig. 1F, Extended Data Fig. 2A**), we did find that group 2 clones were more sensitive to transaminase inhibition (**Fig. 1H, Extended Data Fig. 2b**). Accordingly, as we hypothesized that glutamine was used to fuel mitochondrial metabolism, we tested if this differential sensitivity would extend to inhibition of mitochondrial respiration. Indeed, we observed that group 2 clones were more sensitive to inhibition of mitochondrial respiration through treatment with either the ATP synthase inhibitor oligomycin or the complex I inhibitor phenformin (**Fig 1H,I, Extended Data Fig. 2C**).

### Co-cultures of Different Clonal Populations Provide Metabolic Support

As there was clear distinction among the groups, as it related to baseline metabolism and sensitivity to inhibition of anabolic pathways, we next sought to determine if less sensitive clones would be able to provide metabolic support in the presence of metabolic inhibitors. To accomplish this, we fluorescently labeled an oligomycin sensitive group 2 clone and co-cultured it with either an unlabeled version of itself, a different oligomycin sensitive group 2 clone, or with oligomycin resistant group 1 clones (**Fig. 2A**). Indeed, we observed that the group 1 clones were able to support the growth of the group 2 clones in the presence of a concentration of oligomycin sufficient to block their proliferation when cultured alone or with a different group 2 clone (**Fig. 2B,C**). We further observed that these observations held true with the use of phenformin to inhibit mitochondrial metabolism (**Fig. 2D, Extended Data Fig. 3A,B**). However, the proliferation of a group 1 clone was not further supported in co-culture with a different group 1 clone and grew slightly worse with group 2 clones (**Extended Data Fig. 4**). To validate that this observation was not confined to this set of clonal PDA cells, we obtained a second set of murine KPC clones. Here again we observed differences in oligomycin sensitivity, differential lactate production, and the rescue of mitochondrial inhibition in co-cultures (**Extended Data Fig. 5**). Finally, we observed that the ability of group 1 cells to promote the growth of sensitive group 2 cells in the presence of oligomycin was not contact dependent through transwell co-cultures (**Fig. 2E, Extended Data Fig. 6**).

**Figure 2:**
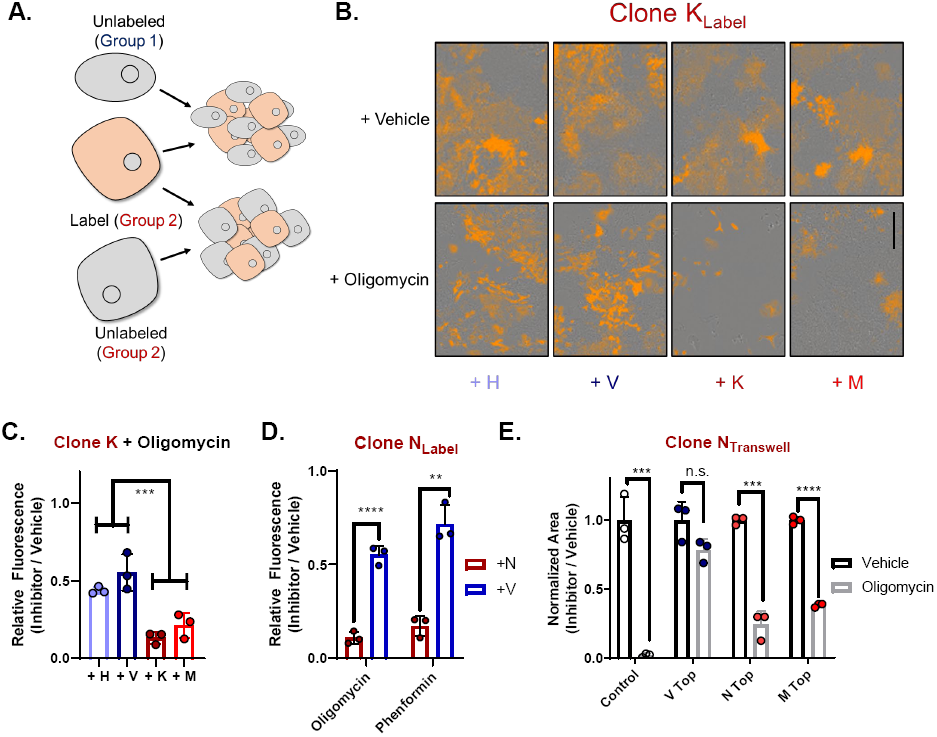
Co-cultures Rescue Mitochondrial Inhibition of Sensitive Clones. **A.** Clonal line K was encoded with a fluorescent label and plated in direct co-cultures with unlabeled clones then treated with oligomycin or vehicle. **B.** Representative images of a labeled oligomycin sensitive clone (K) co-cultured with unlabeled resistant clones (H,V), or with unlabeled sensitive clones (K,M,N) treated with oligomycin or vehicle. **C.** Fluorescent area of oligomycin-treated (0.75 nM) vs. vehicle-treated K-labeled co-cultures from B (n=3). **D.** Fluorescent area of 1 nM oligomycin or 25 µM phenformin treated vs. vehicle for labeled sensitive clone N co-cultured with unlabeled clone N or insensitive clone V (n=3). **E**. Quantitation of colony area of 0.25 nM oligomycin or vehicle treated sensitive clone N with or without transwell inserts containing insensitive clone V, or sensitive clones N, M. Scale bar = 400µm. Error bars are mean ±SD, ** *P* ≤ 0.01; *** *P* ≤ 0.001; **** *P* ≤ 0.0001.

### Asparagine Supports Proliferation During Inhibition of Respiration

We hypothesized that the rescue of proliferation following inhibition of mitochondrial respiration may be mediated through metabolite exchange between the group 1 and group 2 clones. To investigate this, we profiled the media from each of the clonal cell lines by LC/MS-based metabolomics and compared it to their base media. Unsupervised clustering segmented the seven clonal lines into the same two groups as that observed upon analysis of the intercellular metabolite pools (**Extended Data Fig. 7**). Importantly, there was a clear difference in the levels of consumption and release of several metabolites between these groups, largely represented by amino acids and nucleosides, including the production of several non-essential amino acids (NEAAs) (**Fig. 3A,B**).

**Figure 3:**
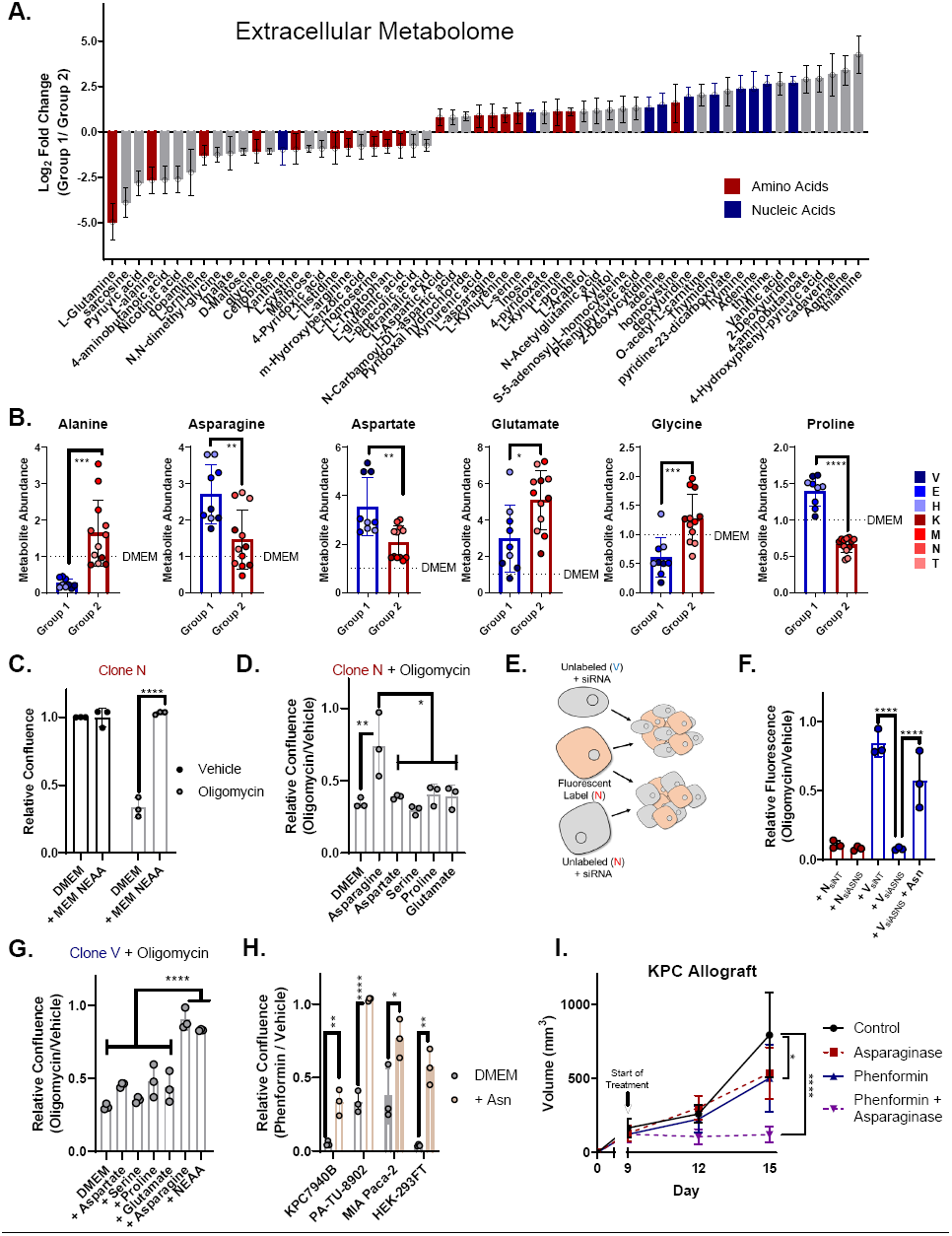
Media Profiling Reveals Asparagine Rescues Inhibition of Respiration. **A.** Differentially consumed/released metabolites present in media from group 1 and group 2 clones (n=3 replicates per cell line) after 48 hours of culture. **B**. Relative abundance of metabolites of non-essential amino acids (NEAAs) present in conditioned media at higher levels than in basal media (n=3 replicates per cell line). **C.** Relative viability of clone N treated with 1nM oligomycin in the presence of absence of media containing all NEAAs (n=3). **D.** Relative viability of clone N treated with 1nM oligomycin in the presence or absence of individual NEAAs (n=3). **E.** Clonal line N was encoded with a fluorescent label and plated in direct co-cultures with unlabeled clones transfect with siRNA targeting *Asns* or non-targeted (NT) control. **F**. Fluorescent area of 1 nM oligomycin treated co-cultures of labeled clone N with unlabeled clone N or insensitive clone V (n=3) transfected with the indicated siRNA with or without the addition of exogenous asparagine. **G.** Relative viability of insensitive clone V treated with 15nM oligomycin in the presence of absence of media containing all NEAAs or individual NEAAs (n=3). **H.** Relative viability of cell lines treated with phenformin with or without exogenous asparagine; KPC7940B and HEK-293FT 150 µM, PA-TU-8902 and MIA PaCa-2 37.5 µM (n=3) **I.** C57B/6J mice were implanted with syngeneic murine PDA cells, and tumors were allowed to establish for 9 days then treated with asparaginase, phenformin, asparaginase + phenformin, or vehicle until harvest. Tumor volume of subcutaneous PDA tumors (n=10). Error bars are mean ±SD, ** *P* ≤ 0.01; *** *P* ≤ 0.001; **** *P* ≤ 0.0001.

Given these observations, we hypothesized that one or more of these amino acids may function to support metabolism in the presence of an inhibitor of mitochondrial respiration. Indeed, we found that treatment of a cocktail of NEAAs at 100µM, based on the physiological levels of several NEAAs measured in pancreatic tumor interstitial fluid^40^, was sufficient to promote growth in the presence of oligomycin (**Fig. 3C**). While aspartate has been identified as a limiting metabolite upon inhibition of mitochondrial metabolism^16,18,41^, we found that among the NEAAs, only asparagine was sufficient to rescue growth in the presence of mitochondrial inhibition (**Fig. 3D**) at this concentration.

To validate asparagine mediates the rescue of proliferation of sensitive clones in co-cultures, we targeted asparagine synthetase (*Asns*) (**Fig. 3E, Extended Data Fig. 8**) in resistant clones by RNAi. As expected, a group 2 clone provided no rescue, whereas group 1 clones transfected with the non-targeting siRNA provided a robust rescue. This was completely abolished by silencing *Asns* (**Fig. 3F**). Additionally, we observed that culturing a less sensitive group 1 clone in doses of oligomycin sufficient to limit their proliferation can also be rescued through addition of asparagine (**Fig. 3G**), suggesting that asparagine rescue of respiration is a general phenomenon. This was further enforced by the observation that the ability of asparagine to rescue proliferation in the presence of phenformin was reproducible across a series of human pancreatic cancer cell lines, and in non-cancerous HEK-293FT cells (**Fig. 3H**).

Given our findings that asparagine can mediate cellular proliferation while respiration is inhibited, we hypothesized that the depletion of extracellular asparagine would function to sensitize tumors to mitochondrial inhibition. To test this, we established syngeneic heterogenous tumors in mice and then treated with PEGylated L-asparaginase, phenformin, or the combination (**Fig. 3I**). In line with our hypothesis, we observed that phenformin treatment showed single agent activity in slowing tumor growth and reducing tumor size. This was greatly enhanced by concurrent asparaginase treatment.

### Exogenous Asparagine Maintains Aspartate and Nucleotide Availability When Respiration is Inhibited

To determine how exogenous asparagine mediates rescue of mitochondrial inhibition, we performed targeted metabolomics on cells treated with oligomycin in the presence or absence of asparagine (**Extended Data Fig. 9**). Consistent with previous data using other mitochondrial poisons, oligomycin treatment depleted TCA cycle and associated branching metabolites, including aspartate (**Fig. 4A**). Aspartate is the biosynthetic precursor for both de novo nucleotide biosynthesis and asparagine. In contrast, human and murine cells are not able to perform the reverse reaction converting exogenous asparagine into aspartate. In the asparagine rescue of oligomycin conditions, we found that cells had both more asparagine and more aspartate (**Fig. 4B,C**). This increase in aspartate permitted the maintenance of nucleotide pools necessary for proliferation (**Fig. 4D**).

**Figure 4:**
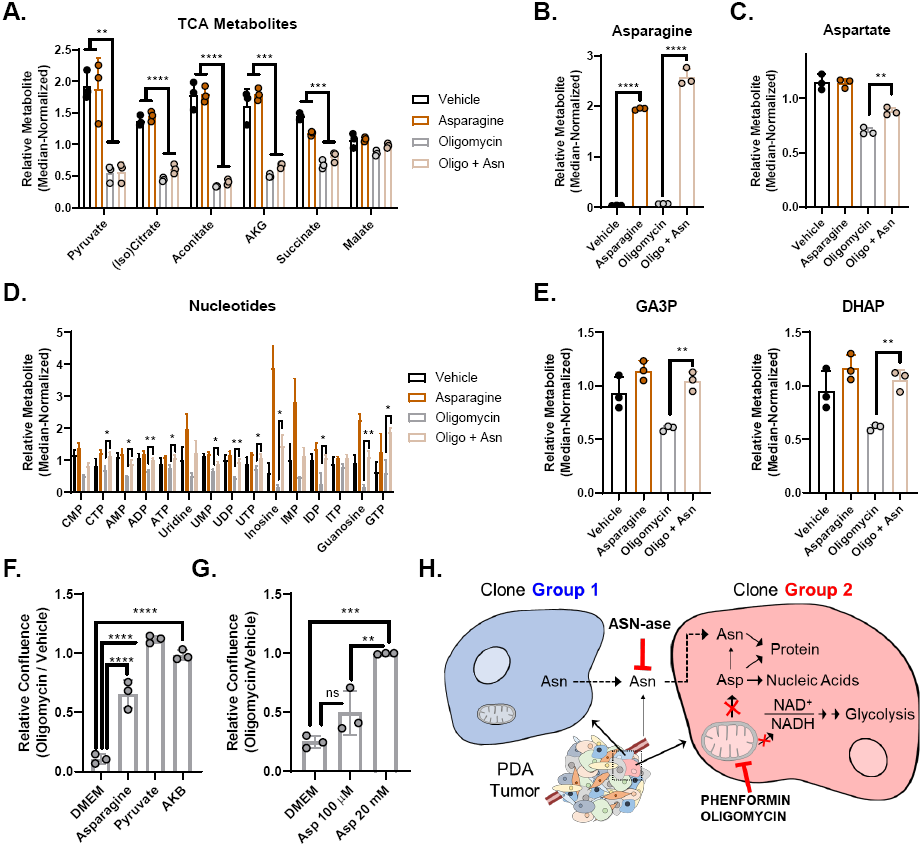
Metabolic Consequences of Exogenous Asparagine During Inhibition of Respiration. **A-E**. Relative abundance of TCA cycle metabolites (**A**), asparagine (**B**), aspartate (**C**), nucleotides (**D**), and glycolytic intermediates (**E**) present in clone N cells treated with asparagine, oligomycin, asparagine + oligomycin, or vehicle 4 hours post treatment (n=3). **F.** Relative viability of clone N treated with 1nM oligomycin in the presence or absence of asparagine, pyruvate, or alpha-ketobutyrate (AKB) (n=3). **G.** Relative viability of clone N treated with 1nM oligomycin in the presence or absence of 100 µM or 20 mM aspartate (n=3). **H.** Group 1 clones in PDA tumors release asparagine (Asn) which can be used by group 2 clones to relieve strain on the aspartate (Asp) pool following inhibition of mitochondrial respiration by oligomycin or phenformin. This allows maintenance of reducing potential, bioenergetics, and precursors for macromolecules required for proliferation. This exchange, and other sources of exogenous asparagine, can be disrupted by asparaginase (ASN-ase) treatment. Error bars are mean ±SD, ** P ≤ 0.01; *** P ≤ 0.001; **** P ≤ 0.0001

Inhibition of respiration also leads to an decrease in the NAD/NADH ratio, which inhibits TCA cycle metabolism, aspartate biosynthesis, and by extension, glycolysis^16,18,41^. We found that, at later time points, asparagine also relieved the NADH-mediated inhibition of metabolism, thereby permitting maintenance of key glycolytic intermediates (**Fig. 4E**). Indeed, treatment with exogenous asparagine rescues oligomycin treatment similar to pyruvate and alpha-ketobutyrate, both of which act to maintain NAD/NADH redox balance in the cell when respiration is impaired (**Fig. 4F**)^18^.

Finally, we found that while aspartate was unable to rescue at micromolar concentrations, like asparagine, it can restore proliferation when provided at 20mM. This difference is likely due to the inability of cells to efficiently uptake aspartate (**Fig. 4G**). Together, these data support the model whereby exogenous asparagine acts, to maintain aspartate, asparagine, and nucleotide pools, which are needed to support proliferation when respiration is impaired (**Fig. 4H**).

## Discussion

Metabolic heterogeneity has been described to result from (i) different mutations in the same tumor type^42^, (ii) the same driving oncogene in different tumor types^43^, (iii) the location of the same tumor cells when seeded at different sites in the body^40^, (iv) regional heterogeneity within the same tumors^44^, and (v) tumor stage (e.g. metastatic versus primary tumor cells)^45,46^. These comparisons illustrate the complexity of cancer metabolism. Furthermore, recent studies in PDA have also illustrated how small subpopulations of cancer cells that survive inhibition of mutant Kras signaling or are capable of anchorage independent growth demonstrate enhanced utilization of mitochondrial oxidative phophorylation^47,48^. Our data build on these collective observations by demonstrating that metabolic complexity is present at baseline within clonal populations derived from the same tumors with a stable genotype.

Despite the considerable potential for metabolic complexity in tumors, our data still demonstrate that it is possible to target metabolism in cancer cells. Indeed, tumor metabolism has been an attractive target in PDA given the limited impact of conventional chemotherapy and immunotherapy on patient survivial^49^. Phenformin was recently identified as the most effective metabolic inhibitor in a screen of PDA patient-derived xenografts^50^, is being explored clinically in other cancer types (NCT03026517), and our data show this can be potentiated by asparagine depletion using asparaginase, a widely used clinically for the treatment of blood cancers^51^. As there are more specific and potent mitochondrial inhibitors currently being deployed in the clinic^52^, including use in pancreatic cancer (NCT03291938), these could potentially also benefit from combination with asparaginase.

Therapeutic application of mitochondrial inhibitors in PDA may have additional avenues of action. For example, we have shown that immune suppressive tumor-associated macrophages in pancreatic cancer primarily utilize mitochondrial metabolism^32^. Accordingly, the use of phenformin and other mitochondrial targeting compounds might also serve to sensitize pancreatic tumors to immune-based therapies, similar to that seen targeting glutamine metabolism and the hexosamine biosynthetic pathway^52,53^. Moreover, efforts to classify targetable growth promoting crosstalk interactions between neoplastic and non-neoplastic cells in the tumor^30-33,35^ are likely to be enhanced by the growing body of data coming out of single cell analyses. Together, these will allow us to continue to find useful new targets for these difficult to treat diseases.

These findings also add to a growing body of literature describing the cellular processes that asparagine mediates to promote proliferation under metabolic stress, such as rescuing apoptosis in the absence of glutamine^54,55^ and mediating mTOR signaling by promoting amino acid exchange^56^. Furthermore, while the role of aspartate in supporting proliferation in the absence of respiration is well established^16,18^, its poor cell permeability appears to require supraphysiological concentrations for aspartate to serve this role in pancreatic cancer. This likely results from absence of an appropriate aspartate transporter, typically restricted to neuronal tissues but seen in some other cancers^17,57^. Given asparagine is capable of promoting proliferation under mitochondrial stress, and at physiological concentrations^40^, there is likely a combination of these identified and further unidentified roles for asparagine, exploration of which will provide exciting new avenues of study.

## Supporting information

combined supplemental submitted

## Materials and Methods

### Cell culture

KPC clonal lines E, H, V, K, M, N, and T were isolated from pancreas tumors from Kras^+/LSLG12D^;Trp^53+/R172H^;Pdx1-Cre (KPC) mice, and have been previously described (https://dx.doi.org/10.2139/ssrn.3486019). Clonal lines 6419c5, 6694c2, 2838c3, and 6499c4 were derived from KPC mice and have been previously described^15^, KPC7940B were a gift from Dr. Gregory Beatty. Cells were maintained in high-glucose DMEM (Gibco) supplemented with 10% FBS (Corning), and routinely tested for mycoplasma contamination using MycoAlert PLUS (Lonza). 2-deoxyglucose, DMSO, oligomycin, phenformin, amnioocyacetic acid were obtained from Sigma.

### Metabolite sample preparation

Intracellular metabolite fractions were prepared from cells grown in 6-well plates (Corning) that were lysed with cold (−80°C) 80% methanol, then clarified by centrifugation. Metabolite levels of intercellular fractions were normalized to the protein content of a parallel sample, and all samples were lyophilized via speed vac after clarification by centrifugation. Media samples were prepared by collecting 200µl of conditioned media or basal media and adding to 800µl of 100% methanol, and the resultant clarified by centrifugation and dried via speed vac. Dried metabolite pellets from cells or media were re-suspended in 35 μL 50:50 HPLC grade MeOH: H2O mixture for metabolomics analysis.

### Metabolomics

Metabolomics were run on either an Agilent 1290 UHPLC-6490 Triple Quadrupole (QqQ) tandem mass spectrometer (MS/MS) system, or an Agilent 1290 Infinity II LC-6470 Triple Quadrupole (QqQ) tandem mass spectrometer (MS/MS) system.

6490 parameters: For negative ion acquisition, a Waters Acquity UPLC BEH amide column (2.1 x 100mm, 1.7µm) was used with the mobile phase (A) consisting of 20 mM ammonium acetate pH 9.6 in water, and mobile phase (B) acetonitrile. The following gradient was used: mobile phase (B) was held at 85% for 1 min, increased to 65% in 12 min, then to 40% in 15 min and held for 5 min before going to initial condition and held for 10 min. For positive ion acquisition, a Waters Acquity UPLC BEH TSS C18 column (2.1 x 100mm, 1.7µm) was used with mobile phase (A) consisting of 0.5 mM NH4F and 0.1% formic acid in water; mobile phase (B) consisting of 0.1% formic acid in acetonitrile. The following gradient was used: mobile phase (B) was held at 1% for 1.5 min, increased to 80% in 15 min, then to 99% in 17 min and held for 2 min before going to initial condition and held for 10 min. The column was kept at 40 °C and 3 µL of sample was injected into the LC-MS/MS with a flow rate of 0.2 mL/min. Tuning and calibration of the QqQ was achieved through Agilent ESI Low Concentration Tuning Mix.

Optimization was performed on the 6490 QqQ in negative or positive mode individually for each of 220 standard compounds to get the best fragment ion and other MS parameters for each standard. Retention time for each standard of the 220 standards was measured from pure standard solution or a mix standard solution. The LC-MS/MS method was created with dynamic (d)MRMs with RTs, RT windows and MRMs of all the 220 standard compounds.

In both acquisition modes, key parameters of AJS ESI were: Gas temp 275 °C, Gas Flow 14 L/min, Nebulizer at 20 psi, Sheath Gas Heater 250 °C, Sheath Gas Flow 11 L/min, Capillary 3000 V. For negative mode MS: Delta EMV was 350 V, Cycle Time 500 ms and Cell accelerator voltage was 4 V, whereas for positive acquisition mode MS: Delta EMV was set at 200 V with no change in cycle time and cell accelerator voltage.

6470 parameters: Agilent Technologies Triple Quad 6470 LC/MS system consists of 1290 Infinity II LC Flexible Pump (Quaternary Pump), 1290 Infinity II Multisampler, 1290 Infinity II Multicolumn Thermostat with 6 port valve and 6470 triple quad mass spectrometer. Agilent Masshunter Workstation Software LC/MS Data Acquisition for 6400 Series Triple Quadrupole MS with Version B.08.02 is used for compound optimization and sample data acquisition.

Solvent A is 97% water and 3% methanol 15 mM acetic acid and 10 mM tributylamine at pH of 5. Solvent C is 15 mM acetic acid and 10 mM tributylamine in methanol. Washing Solvent D is acetonitrile. LC system seal washing solvent 90% water and 10% isopropanol, needle wash solvent 75% methanol, 25% water. GC-grade Tributylamine 99% (ACROS ORGANICS), LC/MS grade acetic acid Optima (Fisher Chemical), InfinityLab Deactivator additive, ESI –L Low concentration Tuning mix (Agilent Technologies), LC-MS grade solvents of water, and acetonitrile, methanol (Millipore), isopropanol (Fisher Chemical).

Agilent ZORBAX RRHD Extend-C18, 2.1 × 150 mm, 1.8 um and ZORBAX Extend Fast Guards for UHPLC are used in the separation. LC gradient profile is: at 0.25 ml/min, 0-2.5 min, 100% A; 7.5 min, 80% A and 20% C; 13 min 55% A and 45% C; 20 min, 1% A and 99% C; 24 min, 1% A and 99% C; 24.05 min, 1% A and 99% D; 27 min, 1% A and 99% D; at 0.8 ml/min, 27.5-31.35 min, 1% A and 99% D; at 0.6 ml/min, 31.50 min, 1% A and 99% D; at 0.4 ml/min, 32.25-39.9 min, 100% A; at 0.25 ml/min, 40 min, 100% A. Column temp is kept at 35 °C, samples are at 4 °C, injection volume is 2 µl.

6470 Triple Quad MS is calibrated with ESI-L Low concentration Tuning mix. Source parameters: Gas temp 150°C, Gas flow 10 l/min, Nebulizer 45 psi, Sheath gas temp 325°C, Sheath gas flow 12 l/min, Capillary −2000 V, Delta EMV −200 V. Dynamic MRM scan type is used with 0.07 min peak width, acquisition time is 24 min. dMRM transitions and other parameters for each compounds are list in a separate sheets. Delta retention time of plus and minus 1 min, fragmentor of 40 eV and cell accelerator of 5 eV are incorporated in the method.

The MassHunter Metabolomics Dynamic MRM Database and Method was used for target identification. Key parameters of AJS ESI were: Gas Temp: 150°C, Gas Flow 13 l/min, Nebulizer 45 psi, Sheath Gas Temp 325°C, Sheath Gas Flow 12 l/min, Capillary 2000 V, Nozzle 500 V. Detector Delta EMV(-) 200.

The QqQ data were pre-processed with Agilent MassHunter Workstation Quantitative Analysis Software (B0700). Additional analyses were post-processed for further quality control in the programming language R. Each sample was normalized by the total intensity of all metabolites to reflect the same protein content as a normalization factor. Finally, each metabolite abundance level in each sample was divided by the median of all abundance levels across all samples for proper comparisons, statistical analyses, and visualizations among metabolites. The statistical significance test was done by a two-tailed t-test with a significance threshold level of 0.05.

Heatmaps were generated and data clustered using Morpheus Matrix Visualization and analysis tool (https://software.broadinstitute.org/morpheus).

### Lactate production measurement

Lactate measurements were carried out using the lactate fluorescence assay kit (Biovision #K607). Assays were performed according to the manufacturer’s instructions. Lactate levels were measured using a SpectraMax M3 Microplate reader (Molecular Devices).

### Western Blotting

Lysates were quantified by BCA assay (Thermo Fisher Scientific) and equal protein amounts were run onto SDS-PAGE gels. Proteins were transferred from SDS-PAGE gel to Immobilon-FL PVDF membrane, blocked, then incubated with primary antibodies. After washing, membranes were then incubated in secondary antibody, washed, then exposed on a Biorad Chemidoc with West PICO ECL (Thermo Fisher Scientific),

### Cell Viability Assay

1000 cells were seeded per well in a 96-well plate and placed on an orbital rocker for 20 minutes to ensure even spreading. Cells were allows to equilibrate overnight, then compounds were added the next day. 2-DG, Oligomycin, AOA, and Phenformin plates were read four days later on the IncuCyte S3 using phase object confluence as a readout. The IC_50_ of 2-DG, oligomycin, and AOA were determined using GraphPad Prism 8. Phenformin sensitivity was determined empirically and data presented by normalizing endpoint confluence to vehicle-treated cells.

### Amino Acid/Pyruvate/AKB Rescues

Cells were treated in triplicate with either a cocktail containing of all NEAAs at 100 µM (Thermo Fisher), or individual amino acids added for a final concentration of 100 µM (Thermo Fisher). 20mM aspartate was prepared directly in DMEM. Pyruvate (Thermo Fisher) was used at 1mM and alpha-ketobutyrate at 500µM. One hour after treatment, oligomycin or vehicle was added. Plates were read four days later on the IncuCyte S3 using phase object confluence as a readout. Sensitivity was determined by treating each cell line at the same concentration and normalizing endpoint confluence to vehicle-treated cells.

### Co-Culture Assays

Cells were grown on clear 96-well plates (Thermo Fisher 167425) at 1000 total cells/well in triplicate and grown for four days. Each well was seeded with 500 fluorescently labeled cells and 500 non-labeled cells, then immediately placed on an orbital rocker for 20 minutes to ensure adequate distribution of cells. Cells were left to in the incubator to attach overnight, and inhibitors were added the following day. At endpoint, total fluorescent area per well was read using the InucCyte S3.

### Asparagine synthetase knock-down

ON-TARGETplus siRNA targeting murine ASNS was purchased from Dharmacon (L-047839-01-0005) with ON-TARGETplus Non-targeting siRNAs as a control (D-001810-10). Cell lines were transfected in 6-well plates using Lipofectamine RNAiMAX Transfection Reagent (Thermo Fisher) per the manufacturer’s instructions. Co-culture viability assays were performed on the IncuCyte S3 as described. Parallel transfected and non-transfected cell lines were harvested for protein to confirm ASNS knockdown via western blot.

### Generation of Fluorescent Cell Lines

Lentiviral constructs containing fluorescent probe were generously donated from Essen BioScience Inc. Cells were seeded at 250,000 cells/well in clear 6-well plates (Corning 3516) and left to attach overnight. The next day, lentiviral constructs were added with polybrene transfection reagent (Sigma). Media was changed 24 hours later.

After 48 hours, puromycin was added to select transfected cells. New media with puromycin was refreshed every 48 hours. Selection was monitored on the IncuCyte until all cells were labelled.

### Antibodies

The following antibodies were used in this study: Vinculin (Cell Signaling 13901), ASNS (Proteintech 14681-1-AP), and Secondary anti-rabbit-HRP (Cell Signaling 7074).

### Transwell Colony-Forming Assay

Cells visualized for colonies were grown on clear 24-well plates (Corning CLS3527) at 200 cells/well for ten days. Cells were grow on inserts (6.5 mm, 4 µm pore size, polyester) at 1000 cells/insert (Corning 3470). 0.25 nM oligomycin or vehicle was refreshed every 48 hours. Colonies were fixed with 100% methanol, stained with (0.5%) crystal violet, and rinsed 6 times with water. Colonies were imaged with Biorad Chemidoc and area quantified using ImageJ.

### Syngeneic Tumor Model

2×10^6^ KPC7940 cells were suspended in a 50:50 matrigel:DMEM mixture and injected subcutaneously in the flanks of syngeneic C57BL/6J mice (Jackson Laboratories). Mice were maintained in SPF housing with access to standard diet and water ad libitum at constant ambient temperature and a 12-hour light cycle. Female mice 8 weeks of age were used for tumor implantation experiments and randomized onto treatment arms, and all experiments were conducted in accordance with the Office of Laboratory Animal Welfare and approved by the Institutional Animal Care and Use Committees of the University of Michigan. Animal numbers were determined in previous work. Phenformin was administered in drinking water containing 5mg/mL sucralose as previously described^58^, and PEGylated asparaginase (Oncaspar) was injected IP at 2UI/100µL PBS every 72 hours.

### Statistical Analysis

Statistics were performed using Graph Pad Prism 8 (Graph Pad Software Inc). Groups of 2 were analyzed with two-tailed students t test, groups greater than 2 were compared using one-way ANOVA analysis with Tukey post hoc test. All error bars represent mean with standard deviation, all group numbers and explanation of significant values are presented within the figure legends. * P ≤ 0.05; ** P ≤ 0.01; *** P ≤ 0.001; **** P ≤ 0.0001.

## Acknowledgements

The authors would like to thank Devon Pendlebury and members of the Lyssiotis lab for scientific feedback. CJH was supported by F32CA228328, K99CA241357, and P30DK034933; BZS was supported by R01CA229803; MPM by the Cancer Moonshot Initiative (U01CA-224145), R01CA151588, and R01CA198074; CJ by a Cancer Research UK Institute Award (A19258) and Experimental Medicine Programme Award (A25236); CAL by a 2017 AACR NextGen Grant for Transformative Cancer Research (17-20-01-LYSS), an ACS Research Scholar Grant (RSG-18-186-01), 1R37CA237421, and R01CA248160; CAL and MPM by the UMCCC Core Grant (P30CA046592). Metabolomics studies were supported by DK097153, the Charles Woodson Research Fund, and the UM Pediatric Brain Tumor Initiative.

## Author Contributions

CJH, CAL conceived of and designed this study. CJH and CAL planned and guided the research and wrote the manuscript. CJH, GT, AM, BSN, PS, AK, PM, LZ, SB, AV, BZS, HC, JZ, and CJ provided key reagents, performed experiments, analyzed, and interpreted data. CAL supervised the work carried out in this study.

