## Supplementary material for "Clonal Heterogeneity Supports Mitochondrial Metabolism in Pancreatic Cancer": combined supplemental submitted

### Extended Data Figure 1: Enriched Metabolic Pathways Among Clonal Metabolites

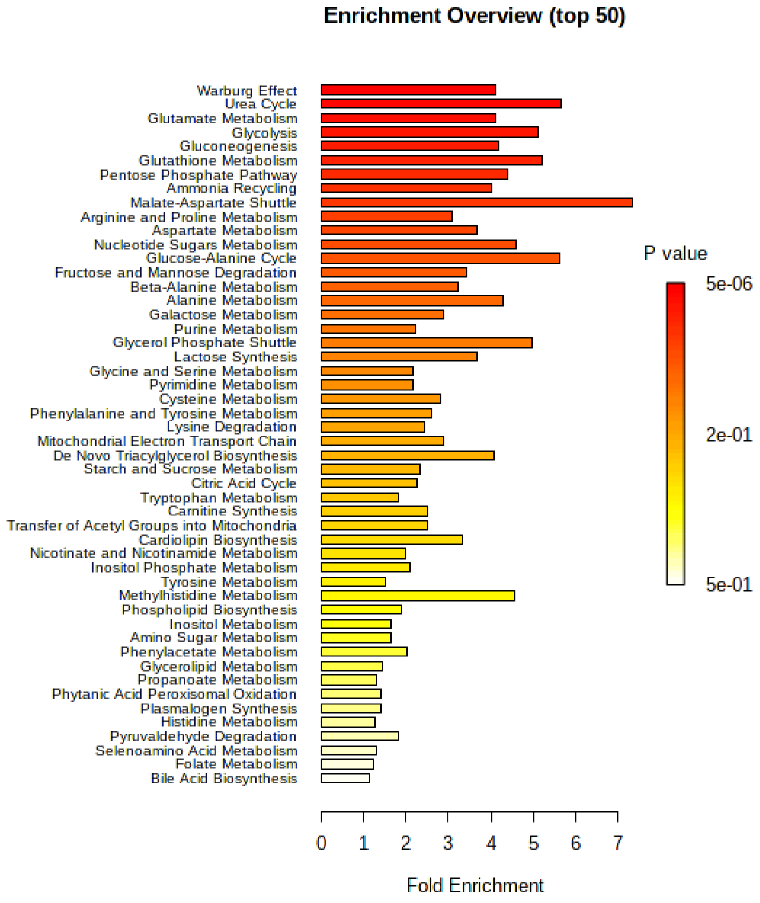

**Extended Data Figure 1:** *Enriched Metabolic Pathways Among Clonal Metabolites.* Top 50 pathways identified by MetaboAnalyst among metabolites differentially represented between group 1 and group 2 clones.

### Extended Data Figure 2: Metabolic Inhibitor Assays

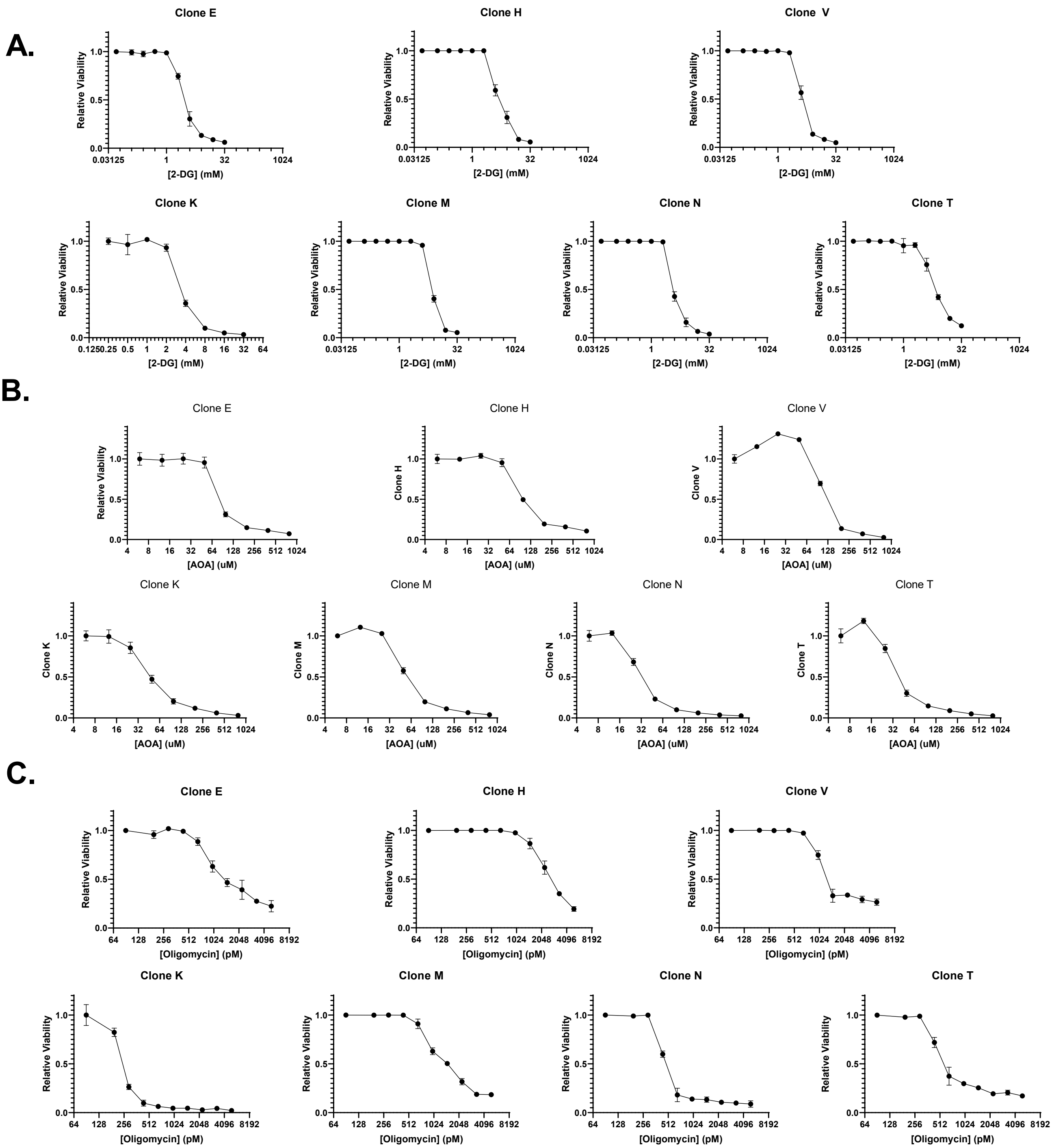

**Extended Data Figure 2:** *Metabolic Inhibitor Assays.* **A.** 2-deoxyglucose (2-DG) dose response curves used to generate $IC_{50}$ values in Figure 1f. **B**. Amnioocyacetic acid (AOA) dose response curves used to generate $IC_{50}$ values in Figure 1g. **C.** Oligomycin dose response curves used to generate $IC_{50}$ values in Figure 1h.

Extended Data Figure 3: Clone N Co-culture Images

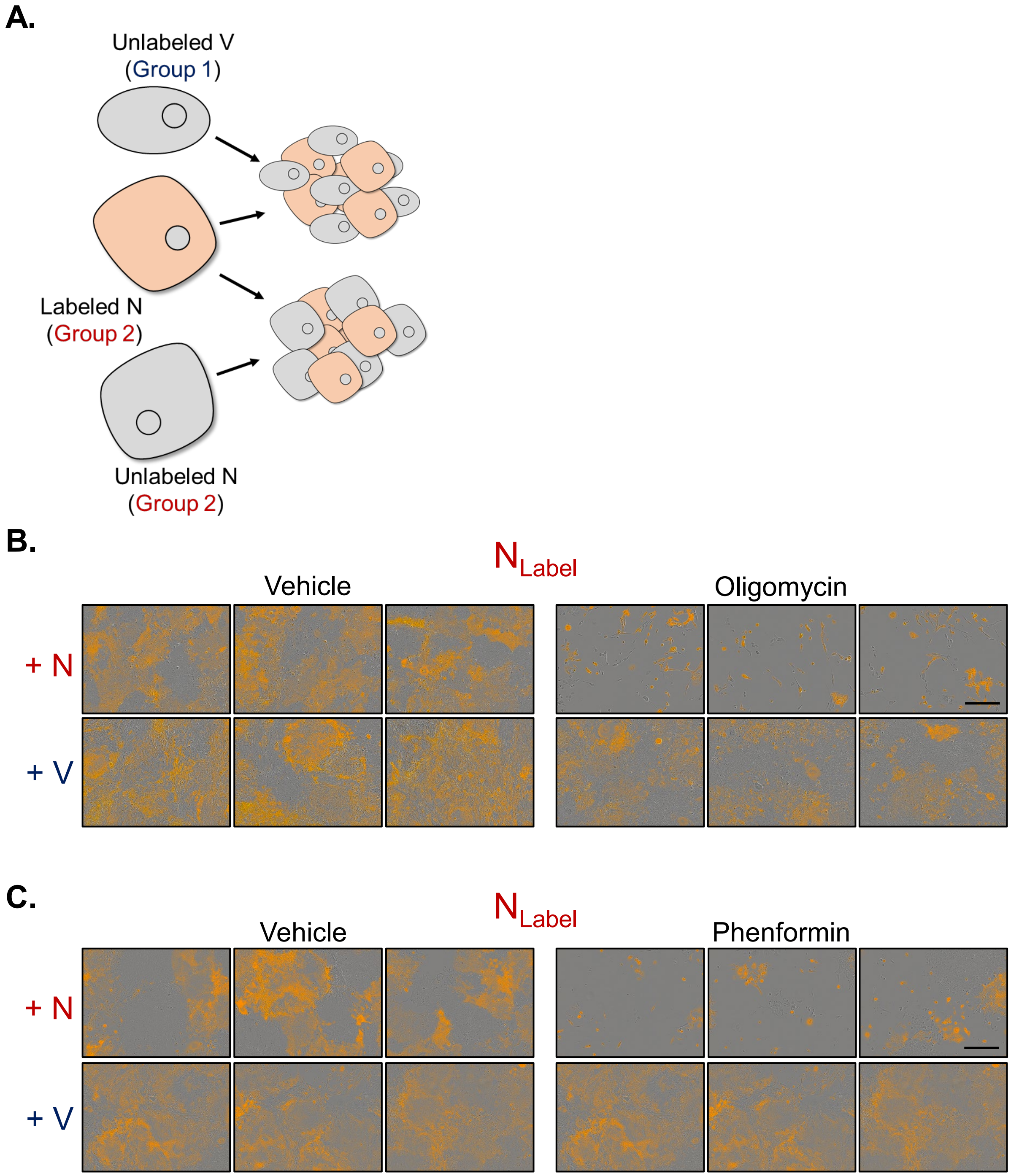

**Extended Data Figure 3:** *Clone N Co-culture Images.* **A.** Group 2 clone N was labeled with a fluorescent probe and co-cultured with either unlabeled N or with unlabeled group 1 clone V and then treated with either phenformin, oligomycin, or vehicle. **B.** Images of 1 nM oligomycin or vehicle treated co-cultures used to generate data presented in Figure 2d. **C.** Images of 25 μM phenformin or vehicle treated co-cultures used to generate data presented in Figure 2d. Scale bar = 400μm

Extended Data Figure 4: Labeled Group 1 Clone Co-cultures

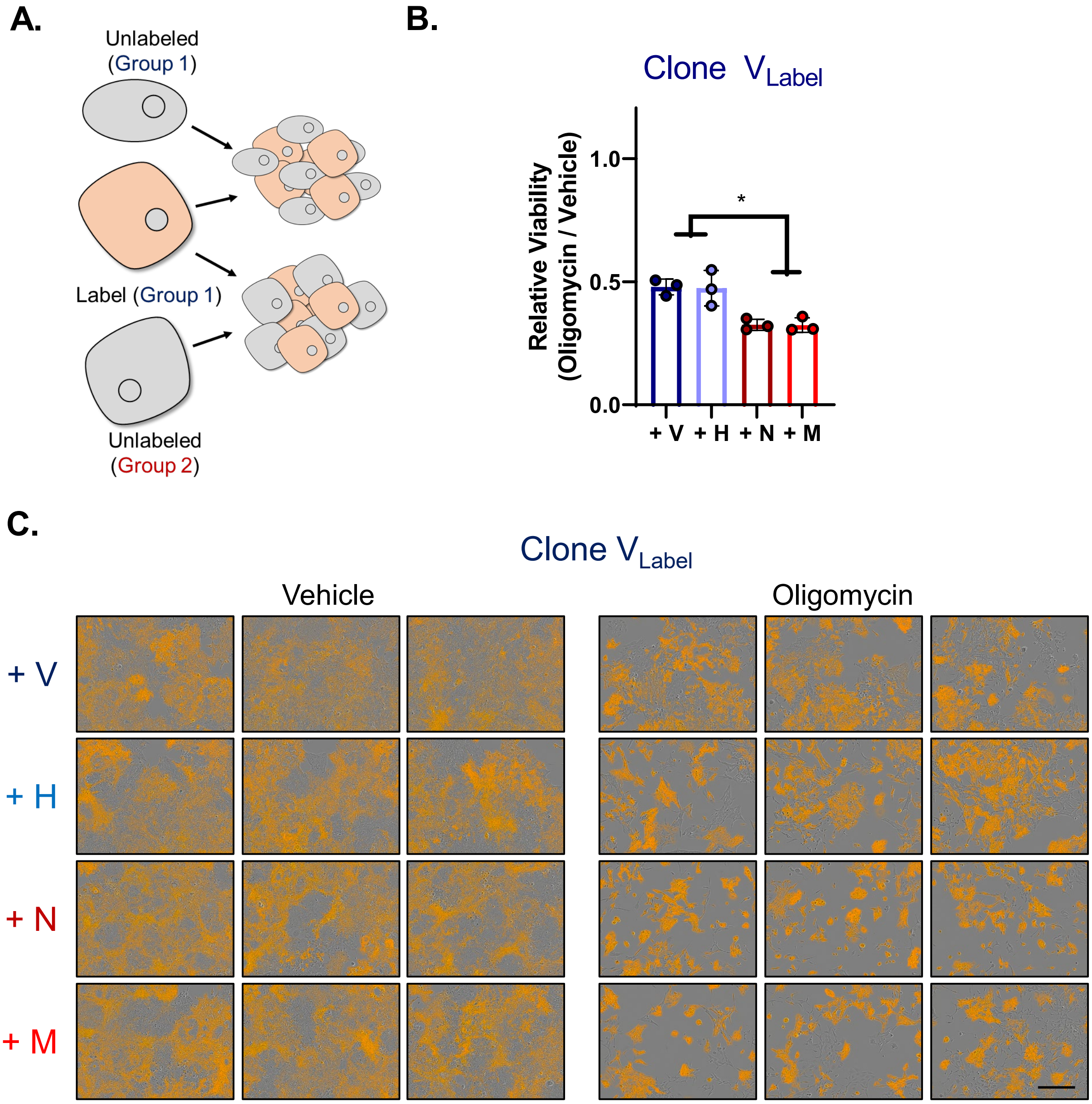

**Extended Data Figure 4**: *Labeled Group 1 Clone Co-cultures*. Group 1 clone V was labeled with a fluorescent probe and co-cultured with either unlabeled group 1 clones (V,H) or with unlabeled group 2 clones (N,M) and then treated with either oligomycin, or vehicle. **B.** Viability of 15 nM oligomycin treated co-cultures relative to vehicle, with representative images (**C**). Error bars are mean ±SD, * $P \leq 0.05$. Scale bar = 400μm

### Extended Data Figure 5. Reproduction in Second Clonal Series

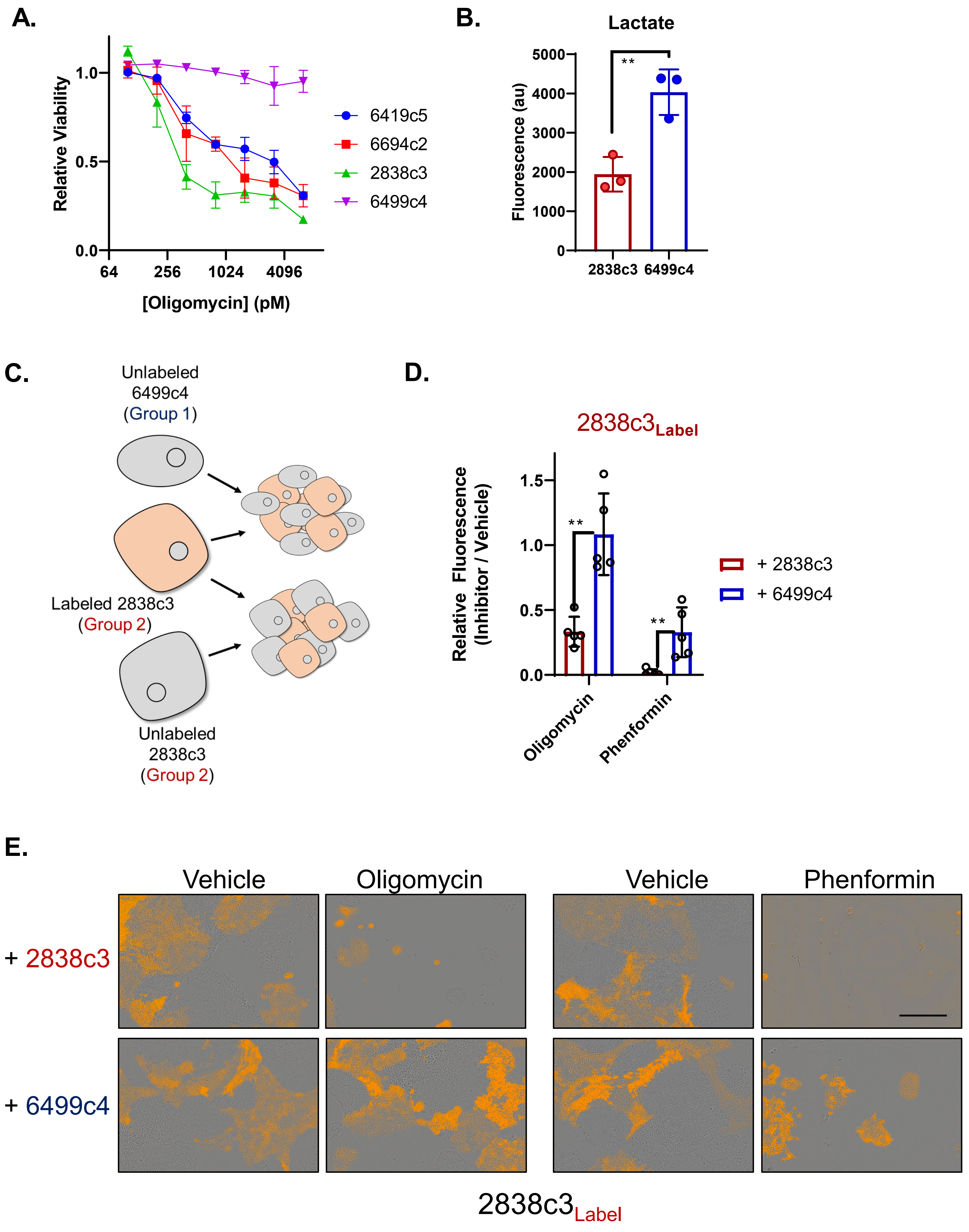

**Extended Data Figure 5:** *Reproduction in Second Clonal Series.* **A.** Dose response curves of oligomycin treated KPC clones 2838C3, 6499c4, 6419c5, and 6694c2. **B.** Lactate production by clones 2838c3 and 6499c4. **C.** Group 2 clone 2838c3 was labeled with a fluorescent probe and co-cultured with either unlabeled 2838c3 or with unlabeled group 1 clone 6499c4 and then treated with either phenformin, oligomycin, or vehicle. **D**. Viability of 5 nM oligomycin or 100 μM phenformin treated co-cultures relative vehicle, with representative images (**E**). Error bars are mean ±SD, ** $P \leq 0.01$. Scale bar = 400μm

### Extended Data Figure 6. Transwell Co-culture Data

**A.**

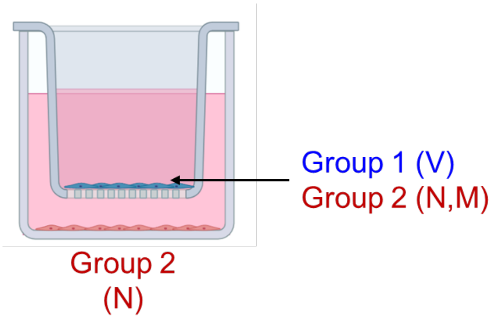


**B.**

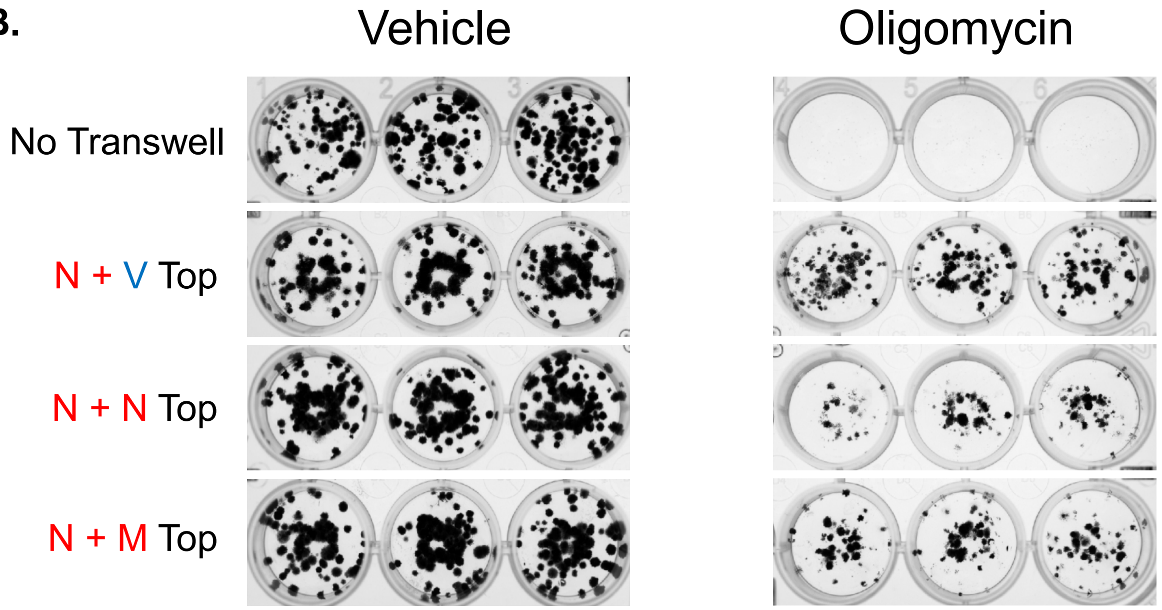


**C.**

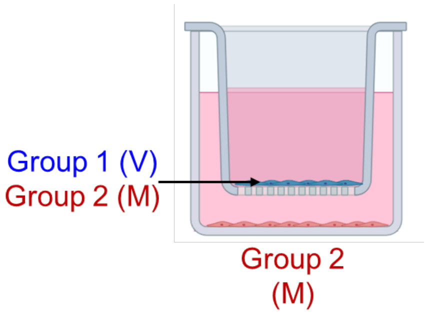


**D.**

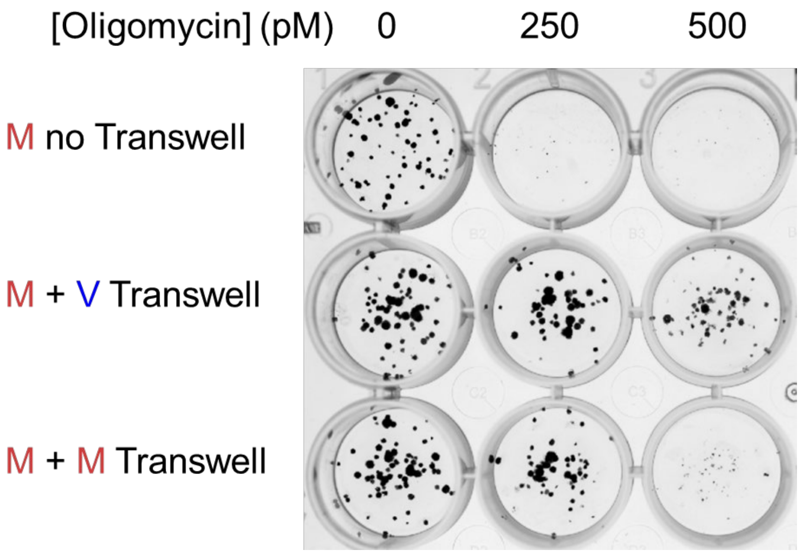

**Extended Data Figure 6**: *Transwell Co-culture Data*. **A.** Group 2 clone N was plated with no transwell, transwells containing group 1 clone V, or group 2 clones (N,M). **B.** Cultures were treated with 0.25 nM oligomycin with fresh drug added in every 48 hours for ten days, then fixed and stained with crystal violet. **C.** Group 2 clone M was plated with no transwell, transwells containing group 1 clone V, or group 2 clones (M). **D.** Cultures were treated with the indicated dose of oligomycin with fresh drug added in every 48 hours for ten days, then fixed and stained with crystal violet.

### Extended Data Figure 7. Significant Media Metabolites

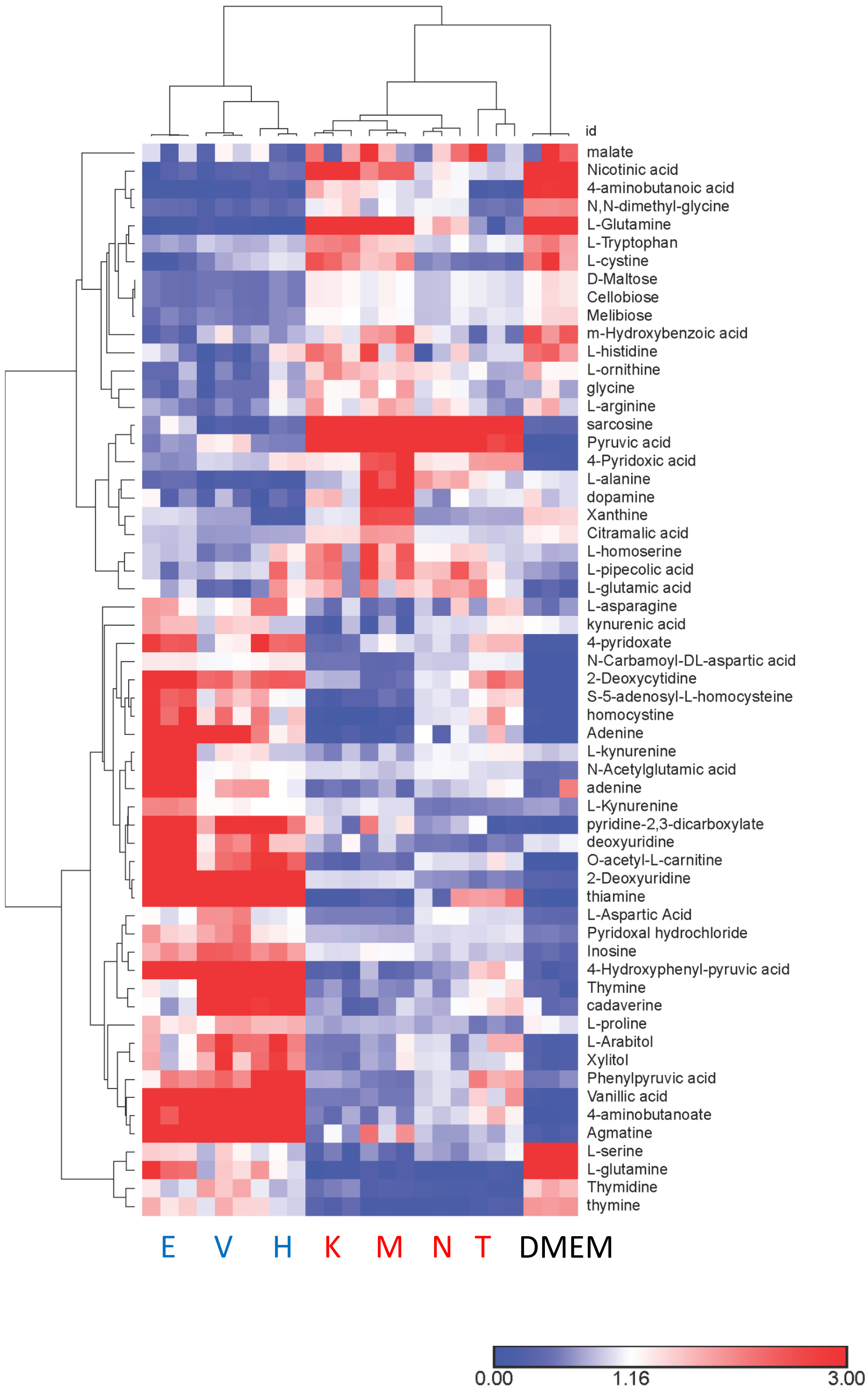

**Extended Data Figure 7:** *Significant Media Metabolites*. Heatmap representation of clustered metabolites from conditioned media after 48 hours of culture which differentiate ±2 fold difference, p=0.01 group 1 clones (E,V,H) vs. group 2 (K,M,N,T).

### Extended Data Figure 8. Asparagine Synthetase Silencing Data

**A.**

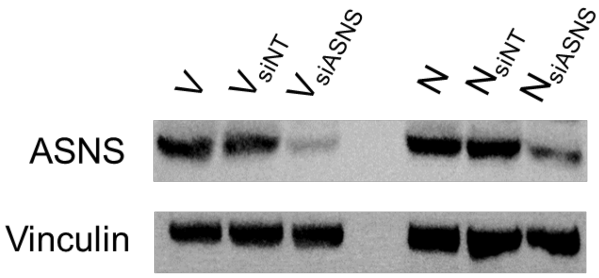


**B.**

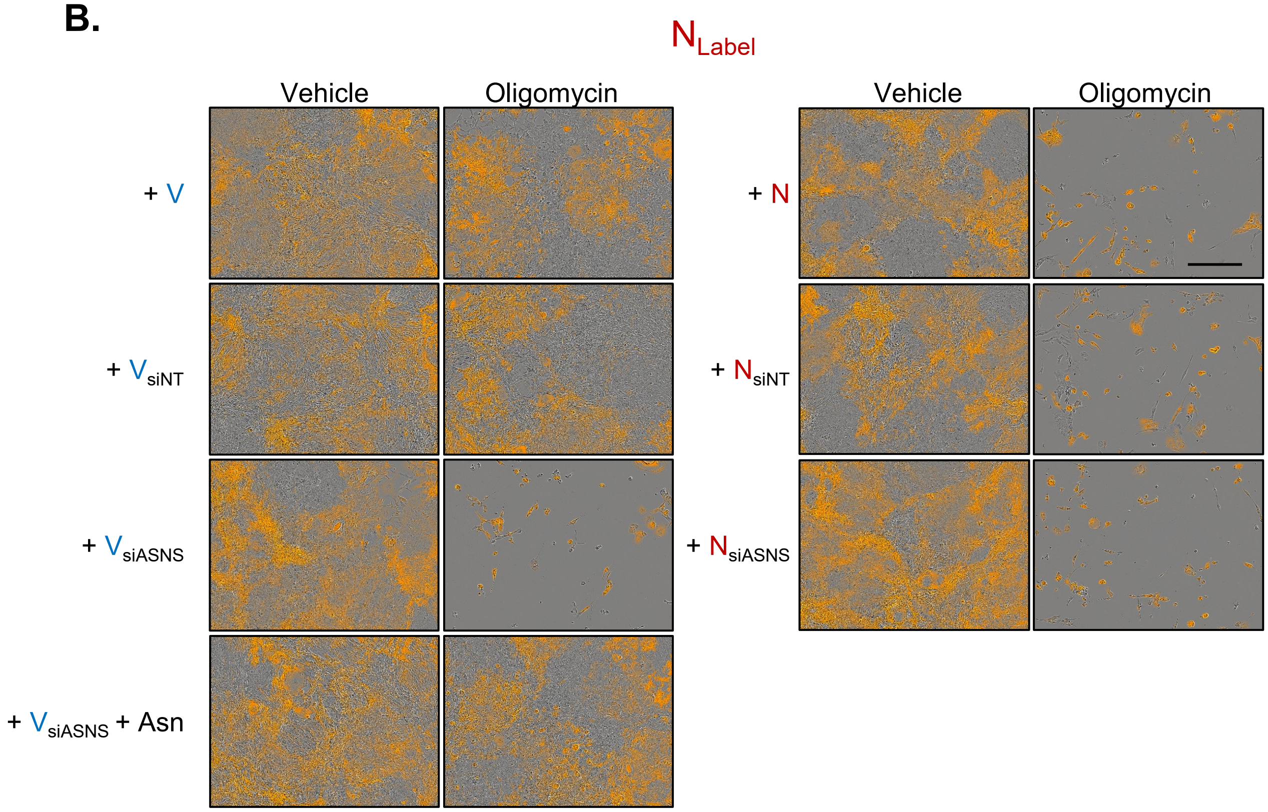

**Extended Data Figure 8:** *Asparagine Synthetase Silencing Data*. **A.** Immunoblot for asparagine synthetase (ASNS) and vinculin of lysates from clones N and V, or clones N and V transfected with non-targeting siRNAs or siRNA targeting *Asns*. **B.** Representative images of 1 nM oligomycin or vehicle treated co-cultures used to generate data presented in Figure 3f. Scale bar = 400µm

### Extended Data Figure 9. Oligomycin + Asn Treatment Significant Metabolites

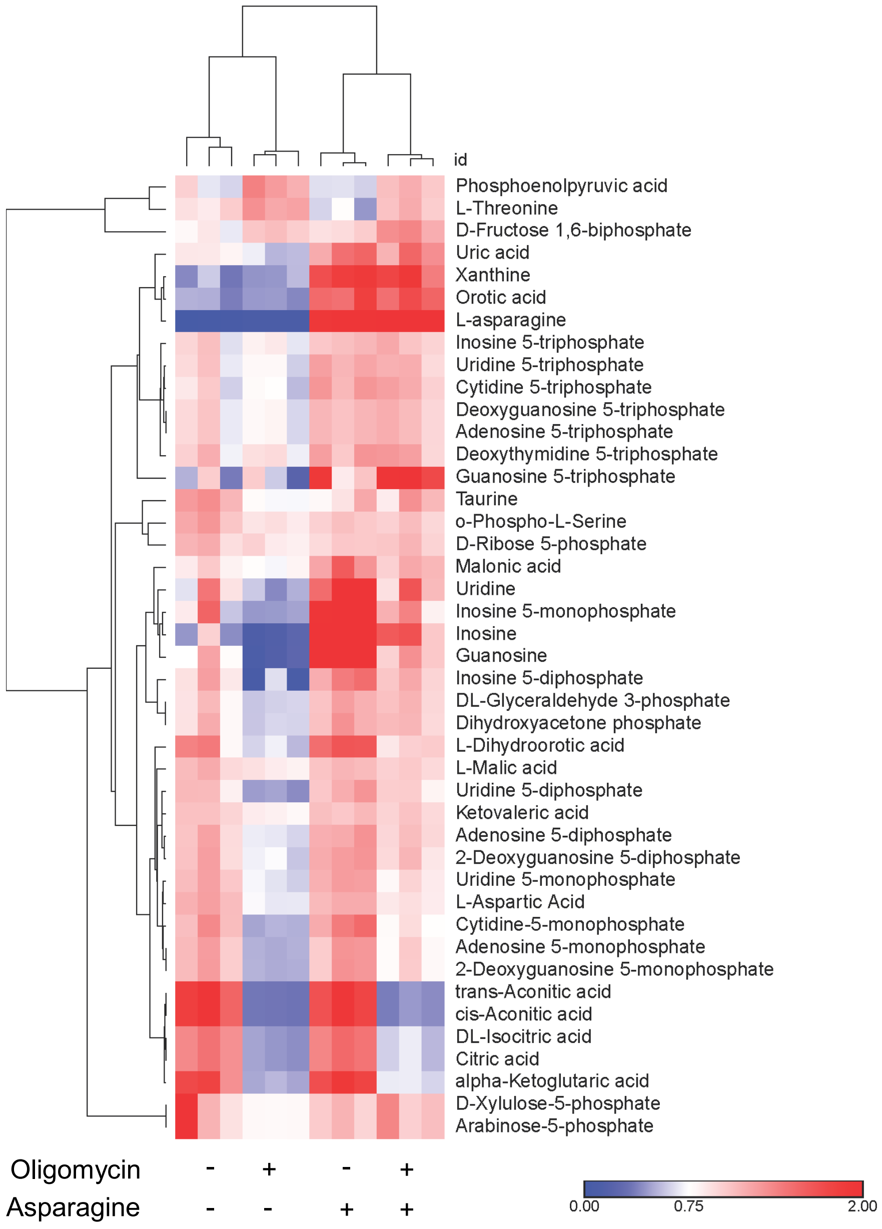

**Extended Data Figure 9:** *Oligomycin + Asn Treatment Significant Metabolites*. Heatmap representation of clustered metabolites ±2 fold difference, p=0.01 from Clone N treated with vehicle, oligomycin, asparagine, or oligomycin + asparagine for 4 hours.
